# Characteristics of a Block to Entry via an HIV-2 Envelope, MCR

**DOI:** 10.1101/177808

**Authors:** Eleanor R. Gray

**Author notes:** Corresponding author. Current address: London Centre for Nanotechnology, University College London, 17-19 Gordon St, London WC1H 0AH.

## Abstract

Successful viral infection depends not only on targeting the correct cell type, but also on the route taken into the cell. After entry, a retrovirus must reverse transcribe its genome and access the nucleus. Blocks to infection can arise at many different stages of the cycle; if they occur only in some cell types it can reveal hitherto unknown aspects of viral or cellular biology. A block to infection of a primary isolate of HIV-2 was previously identified in specific cell types. In this study, parameters of route of entry of the envelope protein from this primary isolate, MCR, were investigated. A critical component of the block acts at a pre-reverse transcription stage, as virions pseudotyped with MCR envelope did not undergo fusion and entry rates commensurate with productive infection. Furthermore, expression of p56^lck^, which regulates CD4 surface expression, partially rescued infection of MCR envelope-pseudotyped virus in restrictive cell types. Based on these findings, we propose that a part of this block results from poor cell-surface expression of CD4.

## Introduction

Innate blocks to retroviral infection of mouse cells were described over 50 years ago (Lilly, 1970, Friend, 1957a,b). In 2000, a capsid (CA)dependent block in other mammalian cell lines was described (Towers et al., 2000) and Trim5*α* was identified in 2004 (Stremlau et al., 2004). Primate variants of Trim5*α* are the causal agent of both HIV-1 restriction in rhesus macaques (Stremlau et al., 2004) and N-MLV restriction in primate cell lines (Keckesova et al., 2004, Yap et al., 2004). Cells resistant to infection under certain circumstances have been discovered both by serendipity and rigorous screens, and this has led to the discovery of novel genes that induce resistance (Gao and Goff, 1999, Bruce et al., 2005, Pineda et al., 2007). Evidence of protective functions of Trims other than Trim5*α* (Yap et al., 2004) has been investigated, as well as variants of Trim5*α* itself in which CypA replaces the B30.2 region (Sayah et al., 2004, Newman et al., 2008, Wilson et al., 2008).

In 2004 a block to replication of HIV-2 by a restriction factor-like agent was described with both capsid (CA) and envelope (Env) dependent components (Schmitz et al., 2004), and this block was named Lv2. Lv2 is dependent on CA sequence, with an additional Env component that can be overcome by pseudotyping with VSV-G. This suggested a route of entry dependence (Schmitz et al., 2004, Marchant et al., 2005). Lv2 was characterised in four cell types; HeLa CD4 cells (Scherer et al., 1953, Maddon et al., 1986) and Ghost/CXCR4 (Cecilia et al., 1998) restrict Lv2-susceptible viruses, whereas NP2/CD4/CXCR4 (NP2) (Soda et al., 1999) and U87/CD4/CXCR4 (U87) (Björndal et al., 1997) do not. Lv2 was initially identified with a primary isolate (prCBL-23) and a T-cell line adapted isolate (CBL-23) of HIV-2, but studies were continued using their equivalent molecular clones, known as MCR and MCN, molecular clone restricted (Lv2 sensitive) and non-restricted (Lv2 insensitive), respectively (Schmitz et al., 2004). These names refer to the whole virus clone of Env and Gag-pol.

The different contributions of Env and CA to Lv2 were described by exchange of MCR and MCN Env with their cognate Gag-pol (Schmitz et al., 2004, Reuter et al., 2005). Virions composed of MCR Gag-pol and Env replicated to over 50-fold higher titres on U87 cells compared to restrictive cells (Schmitz et al., 2004, Reuter et al., 2005), whereas pseudotyping with MCR Env pseudotypes conferred 20-fold restriction on non-susceptible Gag (Schmitz et al., 2004). An additional restriction to infection (of 19- or 5-fold (Schmitz et al., 2004, Reuter et al., 2005)) in HeLa CD4 cells compared to U87 cells was attributable to Gag. Thus Env and Gag act in concert to determine the extent of virus restriction. The determinant in Gag is in CA, at position 207 where there is an isoleucine (MCR) or valine (MCN) (Schmitz et al., 2004). The Env determinant was mapped to position 74 in SU; glutamic acid (E) for MCR or glycine (G) for MCN (Reuter et al., 2005). Lv2 affects HIV-1 cores equivalently to HIV-2 (Marchant et al., 2005, Harrison and McKnight, 2011), showing that Env is a key determinant and can confer sensitivity on an unrelated core. VSV-G pseudotypes of MCR Gag are fully infectious, thus entry via an endocytic route to both Lv2-negative and Lv2-positive cells appears to subvert the block to infection (Schmitz et al., 2004).

If the block to MCR Env-pseudotyped virus is post-entry and route-dependent, then the novel nature of the block is of interest, and could reveal routes of viral entry, and hitherto unknown facets of intracellular compartments. In these studies we describe work to show that conclusive evidence of fusion and entry of restricted virus was not found, hypothesising that the primary cause of the block to infectivity to MCR Env-pseudotyped virus lies in differences in cell biology between restrictive and non-restrictive cells. The cellular basis of the block to MCR Env-pseudotyped virus in HeLa CD4 cells is unlikely to lie within a traditional definition of a restriction factor.

## Methods

### Cells and Viruses

293T, Mus dunni, HeLa CD4 (Scherer et al., 1953), NP2/CD4/CXCR4 (NP2) (Soda et al., 1999) and U87/CD4/CXCR4 (U87) (Björndal et al., 1997) cells were cultivated in DMEM supplemented with 5% (except 293T and Mus dunni, 10%) foetal calf serum and antibiotics. HeLa CD4, NP2 and U87 cells were gifts from Professor Áine McKnight (Queen Mary, University of London). 'Tester' viruses were generated essentially as described in (Bock et al., 2000) with 7*μ*g of the following vectors using a calcium phosphate-based transfection method (Promega): HIV-1 virus-like particles (VLPs) CSGW (Bainbridge et al., 2001) and p8.91 (Naldini et al., 1996); HIV-2 VLPs HIV2 and HIV-2pack (Ylinen et al., 2005); SIV3+ (Besnier et al., 2002); Envelope vectors were MCR (a gift from Professor Áine McKnight), p456 (Soneoka et al., 1995) or pczVSV-G (Bock et al., 2000) to produce MCR-, amphotropic- or VSV-G-pseudotyped VLPs. If S15-mCherry (Campbell et al., 2007) and GFP-Vpr (McDonald et al., 2002) were included, 5*μ*g each of CSGW, p8.91, VSV-G or MCR, and GFP-Vpr were used with 0.5*μ*g S5-mCherry. For generation of p56^*lck*^ vector, the cDNA was obtained from the IMAGE Consortium (Lennon et al., 1996) and cloned into the gateway vector pLgatewayIRESYFP (Yap et al., 2007), and used with pHIT60 (Soneoka et al., 1995) and pczVSVG. 18h post-transfection, cells were treated with 10mM sodium butyrate for 6 h. 48 h after transfection, virus was harvested, filtered through a 0.45*μ*m filter, and stored immediately at −80*°* C. Viral stocks were titrated on Mus dunni cells if pseudotyped with VSV-G or amphotropic envelope, or on NP2 cells if MCR Env. 5*×*10^4^ cells were plated and allowed to settle. 24 hours later, 250-500*μ*l virus stock was added to a well in a total volume of 500*μ*l, and serial 1:5 dilutions made for at least 8 wells. 24 hours later the media was refreshed, and after a further 48 hours titres were assessed by FACS. All viruses used were HIV-1 based, unless specifically stated otherwise.

### Restriction Assays

Single colour assays for entry and a single round of infection were carried out as described previously (Bock et al., 2000). Briefly, 5*×*10^4^ cells were seeded in 12-well plates, challenged after 16hrs with MOI 0.001 to 10 VSV-G, amphotropic MLV envelope (ampho), or MCR Env-pseudotyped virus. 48hrs later, cells were harvested, fixed in phosphate-buffered saline (PBS)-3.5% formalin, and titres assessed by FACS (BD LSRII, default filter set). Dual colour (YFP/GFP) assays for entry and a single round of infection for cells transduced with delivery vector encoding p56^*lck*^ and eYFP were carried out as described previously (Ohkura et al., 2011). Briefly, 5*×*10^4^ cells were seeded in 12-well plates and transduced after 16hrs with MOI 1 p56^*lck*^ vector. After a further 48hrs, cells were trypsinized and reseeded at 1:6 (NP2/U87) or 1:12 (HeLa CD4), then challenged after 16hrs with tester virus. 48hrs later, cells were harvested, fixed in PBS-3.5% formalin, and titres assessed by FACS (BD LSRII with YFP/GFP filter set, namely 550/30 and 525LP for YFP, 510/20 and 505LP for GFP).

### Quantitative PCR

Extraction, quantitation, amplification and normalization were performed essentially as described in (Yap et al., 2006). Briefly, 2*×*10^5^ cells were plated, and 24 hours later incubated with virus at MOI of 2-5 for 6 or 18 hours. Total DNA was extracted with the QIAamp DNA kit (Qiagen). HIV-1 early reverse transcription products were detected using primers HIVEF and HIVER (Yap et al., 2006).

### Microscopy and Particle Counting

Microscopy was performed essentially as described in (Mortuza et al., 2008) with the following specific constraints for sample preparation. Cells were plated on glass coverslips in a 12-well dish at 1*×*10^5^ cells/well, and infected 16 hours later at 4*°* C with virus at MOI 5. Cells were spinoculated at 15*°* C and 1200*×*g for 2 hours, then 1ml pre-warmed DMEM ±40mM NH_4_ Cl was added to each well, and infection allowed to proceed at 37*°* C for 2hrs. To end infection, cells were washed 1 *×* with PBS and incubated with 4% paraformaldehyde in PBS at room temperature for 15 minutes. Slides were viewed on a Deltavision Olympus IX70 inverted microscope with 100x lens and initially analysed using Softworx image acquisition software (Applied Precision). Where necessary, images were further viewed using Adobe Photoshop CS2 (Adobe Systems) which permits removal of each red/green/blue channel individually. For counting of virions, pictures were randomised and renamed by a third party, then scored blind for the extent of overlay of green and red fluorescence.

## Results

### HeLa CD4 and NP2 cells are equally permissive to a selection of HIV-1, HIV-2 and SIV cores pseudotyped with VSV-G

HIV-1, HIV-2 and SIV viral cores were pseudotyped with VSV-G or MCR Env to assess the permissivity of HeLa CD4 and NP2 cells. The HIV-2 vector, derived from ROD10, has an MCR-type isoleucine at Gag 207. Titres of VSVG pseudotyped virus were equivalent for all cores on both cell types (Figure 1a-c). Titres of MCR Env-pseudotyped virus were equally high on NP2 cells, but on HeLa CD4 cells titres were 10 to 100-fold lower. This confirms the route of entry dependence of the Lv2-phenotype, in that the same core pseudotyped with different envelope proteins can exhibit different levels of infectivity.

**Figure 1:**
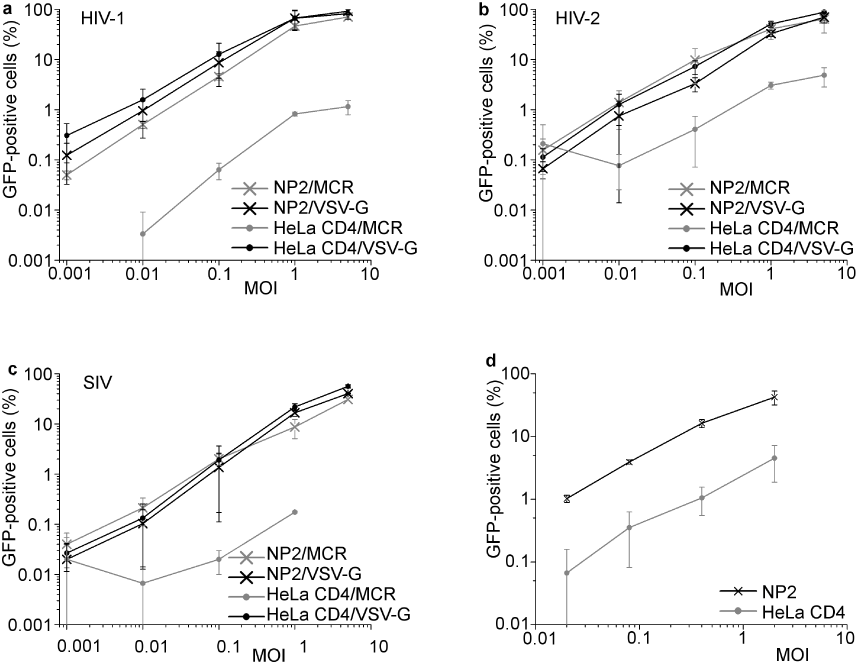
Virus pseudotyped with MCR Env does not successfully replicate on HeLa CD4 cells, but does on NP2 cells. NP2 and HeLa CD4 cells were transduced with MCR and VSV-G pseudotyped a) HIV-1, b) HIV-2 and c) SIVmac cores, and analysed 3 days later by FACS. Viral titres were equivalent for NP2 cells, but MCR Env-pseudotyped virus titres were reduced 10- to 100-fold on HeLa CD4 cells. Shown are the combined results from 3 independent experiments. d) HIV-1 cores pseudotyped with NL4-3 envelope are not able to replicate to high titres on HeLa CD4 cells. Cells were infected with NL4-3pseudotyped HIV-1 core at MOI 0.02 to 2, and analysed 3 days later by FACS. Shown are the combined results of four independent experiments for both cell types, and error bars represent the standard deviation of the mean.

MCR and MCN bind to CD4 and CXCR4 (McKnight et al., 2001). Pseudotyping virions with MCN Env did not circumvent the block to infection seen in HeLa CD4 cells with a considerable 10-100-fold restriction compared to NP2 or U87 cells (Supplementary Figure 1). As MCN Env-pseudotyped virions were thus significantly restricted on HeLa CD4 cells, this could not be used as a control for further experiments. To assess whether another, different CD4/CXCR4-tropic envelope also renders the virus susceptible to this block to infection, NL4-3 Env was used. Figure 1d shows that for all input titres of NL4-3 Env-pseudotyped virus, NP2 cells are between 10to 100-fold more permissive compared to HeLa CD4 cells. The difference was greatest at lower viral titres. Therefore the block to infectivity is not unique to MCR Env, but also affects MCN Env and at least one other CD4/CXCR4-tropic envelope.

### Reduced levels of reverse transcription products are seen in HeLa CD4 cells infected with MCR pseudotyped virus, compared to NP2 cells

Analysis of the first products of reverse transcription reveals whether the virus can enter the cell and reverse transcribe. Figure 2a shows a comparison of relative strong stop transcript levels at 6 and 18hrs of MCR and VSV-G pseudotyped virus on HeLa CD4 cells. Levels of MCR transcripts reach 21% of those of VSV-G (which is set to 100%) at both 6 and 18hr timepoints. Figure 2b shows the level of reverse transcription products in HeLa CD4 and NP2 cells after infection with VSV-G or MCR pseudotyped virus at 18hrs. Infection by virions pseudotyped with these two envelopes produced equivalent levels of strong-stop transcripts in NP2 cells, but not HeLa CD4 cells. For virions pseudotyped with MCR Env, levels of strong stop transcripts in HeLa CD4 cells were 24% of those obtained from an infection with VSV-G pseudotyped virus. These data show that the accumulation of appreciable levels of strong stop transcripts does not occur in HeLaCD4 cells for viruses pseudotyped with MCR Env.

**Figure 2:**
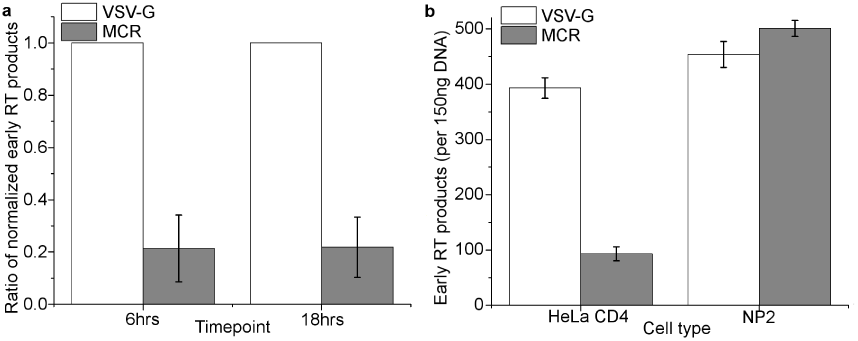
Reverse transcription product levels from HeLa CD4 and NP2 cells infected with virus pseudotyped with VSV-G or MCR Env. Cells were plated out, and 24 hrs later were incubated with virions at MOI 0.8 to 1 at 4° C for 1 hour to permit binding. Infection was initiated by addition of pre-warmed medium, and transfer to 37° C. a) After 6 or 18 hrs cells were harvested, and DNA extracted from HeLa CD4 cells alone. Levels of early products of reverse transcription were assessed by qPCR, and the levels of DNA normalized using actin. Results shown are combined from four independent experiments. b) After 18 hrs cells were harvested and DNA extracted from HeLa CD4 and NP2 cells, levels of early reverse transcription products were again assessed by qPCR and normalized to actin. The level of reverse transcription products for VSVG-pseudotyped viruses is set to 100%. VSV-G data shown in white, and MCR data in grey bars. Results shown are combined from four independent experiments, and error bars represent the standard deviation of the mean.

Two examples of a pre-reverse transcription block caused by a restriction factor are Trim5*α* and Trim5Cyp (Nisole et al., 2004, Stremlau et al., 2004). Infection of Trim5*α* or Trim5Cyp-positive cells with susceptible virus leads to a drop in reverse transcription intermediates that varies between 5 and 10-fold (Yap et al., 2006, 2007, Stremlau et al., 2004, Fletcher et al., 2010, Anderson et al., 2006) (although the difference in infectivity caused by these blocks can be greater than 10-fold) so the drop seen here is commensurate with the lack of significant synthesis of reverse transcription products as seen for other restriction factors.

These results are also comparable to previous studies comparing levels of reverse transcripts by virions pseudotyped with MCR Env produced from HeLa CD4 and unrestrictive cells. The original primary isolate later cloned as MCR, prCBL23, produced higher levels of LTR-gag on U87 cells compared to HeLa CD4 cells (McKnight et al., 2001). Schmitz et al. found ratios of strong-stop reverse transcripts for MCR pseudotyped virions were between 7-25-fold higher from U87 cells compared to HeLa CD4 cells at timepoints between 1 and 18hrs (Schmitz et al., 2004). Marchant et al. also reported greatly reduced levels of reverse transcripts from HeLa CD4 cells compared to U87 cells from MCR Env pseudotyped virus at 2 and 6hrs, though by 18hrs the levels had equalised (Marchant et al., 2005). This suggests that MCR Env-pseudotyped virions are unable to produce levels of reverse transcripts in HeLa CD4 cells that are sufficient for a productive infection.

### The block to infection in HeLa CD4 cells cannot be abrogated

To assess whether the block to MCR-pseudotyped virions can be saturated, in common with Fv1 and Trim5*α*, the susceptibility of the block to abrogation was assessed, as shown in Supplementary Figure 2. No increase in titres was seen when a prior, restricted dose of virus was applied, indicating that the block to infection cannot be saturated (compare to Towers et al. (2002), Dodding et al. (2005)). This means that the block to infection is dissimilar in nature to saturable restriction factors such as Fv1 and Trim5*α*.

### Levels of viral fusion can be monitored using virions produced in cells expressing s15mCherry

MCR Env-pseudotyped virions exhibit lower infectivity on HeLa CD4 cells and do not reverse transcribe at equivalent levels to those on NP2 cells. Therefore the antecedent stage to investigate in the process was fusion, which must occur prior to reverse transcription. A fluorescent fusion protein comprised of mCherry and the 15 N-terminal amino acids of Src (s15-mC) coats nascent viral particles with a red membrane as they leave the producer cell (Rodgers, 2002, Campbell et al., 2007). The signal disperses and is lost after fusion between the viral and cellular membranes has taken place. If fusion is blocked then the red signal remains intact (Campbell et al., 2007). This system was used to analyse whether fusion was successfully taking place between MCR pseudotyped viruses and HeLa CD4 cells. To monitor virions after fusion, the fluorescent core protein GFP-Vpr was used (McDonald et al., 2002). Viral particles outside the cell, and those bound to cells pre-fusion remain both green and red; those that have successfully fused are green only.

For proof of concept, GFP-vpr and s15-mC-containing viral particles were pseudotyped with VSV-G envelope protein, bound to HeLa CD4 or U87 cells on coverslips and then incubated for two hours in the presence or absence of NH_4_ Cl. This blocks initiation of fusion by the VSV-G envelope so that the virus remains trapped in endosomes (Aiken, 1997, Brindley and Maury, 2005, Superti et al., 1987). After infection and fixation, at least 10 pictures were taken, randomised, and scored blind for overlapping red and green, or green alone, punctae. Examples of pictures used to obtain data are shown in Figure 3a, and the percentage of green vs. green and red particles in each of the different conditions quantified in Figure 3b.

**Figure 3:**
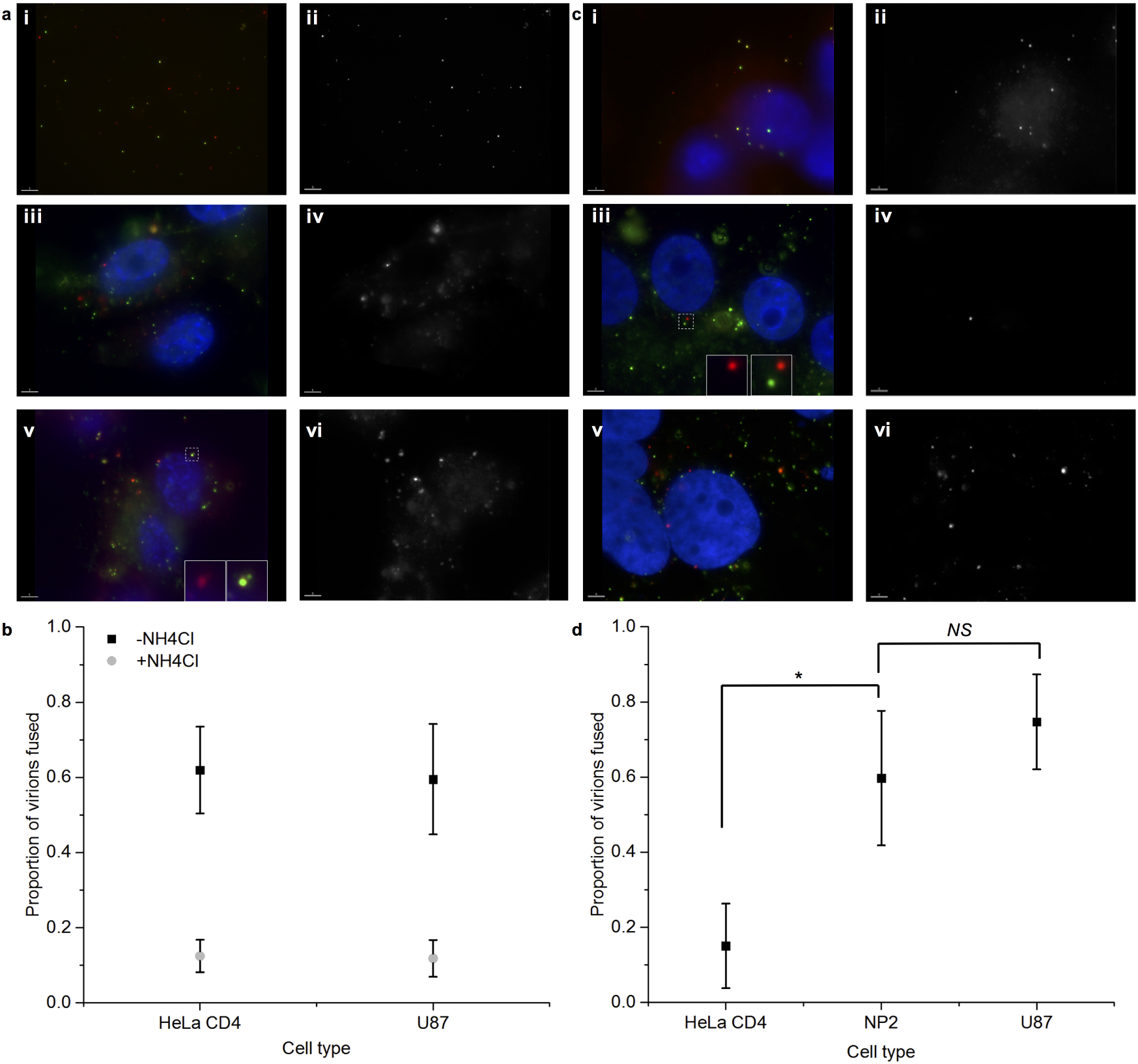
Virions pseudotyped with VSV-G or MCR Env visualised after binding to HeLa CD4, NP2 or U87 cells. Cells were infected with GFP-vpr/S15-mCherry virus pseudotyped with A+B) VSV-G in the presence or absence of NH4Cl, or C+D) MCR Env at MOI 1. Images in (a) and (c) are shown for RGB channels (i, iii and v) or R channel only (in greyscale, ii, iv and vi). a) i+ii) VSV-G pseudotyped virions used to infect cells, showing red and green fluorescence; a) iii to vi) HeLa CD4 cells and virions pseudodyped with VSV-G without (iii + iv) and with (v and vi) NH4Cl. c) virions pseudotyped with MCR Env and (i + ii) HeLa CD4 cells, (iii + iv) NP2 cells and (v + vi) U87 cells. A representative picture of at least 10 taken per condition is shown for (a) and (c). The boxes in (a)(v) and (c)(iii) show two identical enlarged views of the section enclosed by dashed lines, with the green channel removed for the second view to show an example of a fused and unfused virion in each case. The quantified results in (b) and (d) are the proportions of virions that have successfully fused with the cell, lost their red membrane signal and show GFP fluorescence only. Proportions per picture are shown, averaged from at least 10 separate pictures taken per condition, randomised, and over 500 virions per condition counted and scored blind for red or red and green fluorescence. All images were taken at the same scale. Error bars show standard deviation of the mean number of virions fused per picture. Results from one experiment are shown of two performed. *denotes p< 0.05, NS=not significant (Student's t-test).

The data in Figure 3b show that after 2 hours of incubation with media alone, 62 and 59% of viral particles had undergone fusion in HeLa CD4 and U87 cells respectively. If NH_4_ Cl was present, successful fusion events were seen in only 12% of particles tallied, for both cell types. Assured detection of the presence of a red label is harder than its absence, so the percentage of fusion events in cells could be even lower than these values. This method provides an alternative to qPCR to assess whether virions have entered cells, and permits assessment of the post-binding fusion of the viral and cellular membranes at a stage before reverse transcription.

### Levels of fusion between HeLa CD4 cells and MCR pseudotyped HIV-1 are insufficient for infection

The S15-mC and GFP-vpr labels were used to assess the proportion of MCR Env-pseudotyped viral particles that underwent fusion in HeLa CD4, NP2 and U87 cells. Two cell lines permissive to MCR were assessed to types. Sample pictures are shown in Figure 3c and the quantified data in Figure 3d.

A significant difference in the proportion of virions fused was seen between either HeLa CD4 and U87 cells, or HeLa CD4 and NP2 cells. The proportion of MCR virions that underwent fusion per picture was 15%, 60%, and 75% for HeLa CD4, NP2 and U87 cells, respectively (Figure 3d). Furthermore, the difference in fusion levels between HeLa CD4 cells with MCR pseudotyped virus, or HeLa CD4 cells with VSV-G pseudotyped virus held under an NH_4_ Cl-induced block to infection was not significant. This indicates that the level of fusion is insufficient for a productive infection for both conditions.

Interestingly, the average number of MCR virions per picture of HeLa CD4 cells was 21.5, which was lower than for VSV-G virions with HeLa CD4 cells (93.2), or MCR virions with NP2 cells (81.7) and U87 cells (46.2). This is an incidental observation, but could suggest that CD4 binding of MCR Env is easily disrupted, perhaps by the washing steps. Antibody binding studies by Reuter et al., however, showed that MCR Env actually conferred a ten-fold resistance to inhibition of infection by anti-CD4 antibody compared to unrestricted envelope (MCN), concluding that MCR binds to CD4 more tightly (Reuter et al., 2005). So it is more likely that MCR does not readily bind in the first place. Platt et al. suggest that the low affinities for CD4 seen for patient isolates envelopes generally may be an adaptation that allows them to bypass cells that would not result in a productive infection and have low levels of CD4 at the cells surface, such as early T cell progenitors (Platt et al., 2000). Alternatively, the turnover of virions by HeLa CD4 cells could be more rapid, with a higher rate of recycling of bound virus.

These data, showing that MCR Env-pseudotyped virions do not fuse with HeLa CD4 cells at levels sufficient for a productive infection, combined with the lack of products of reverse transcription and non-saturation of the block, would suggest that there could be a difference in presentation, signalling or regulation of these receptors in HeLa CD4 cells that is different to NP2 and U87 cells, and this possibility was next examined.

### Expression of p56^*lck*^ renders HeLa CD4 cells moderately permissive to MCR pseudotyped virions

CD4 is a marker of differentiation found primarily on T cells, and control of CD4 levels at the cell surface is mediated by the tyrosine kinase p56^*lck*^, which is constitutively associated with the cytoplasmic domain, and plays a key role in regulating endocytosis (Pelchen-Matthews et al., 1991, 1992, 1995, Pitcher et al., 1999). CD4 that has no cytoplasmic domain and cannot interact with p56^*lck*^ is likewise constitutively endocytosed (Pelchen-Matthews et al., 1991, 1995, Foti et al., 2002, Yousefi et al., 2003). HeLa cells are not thymocytes, and the presence or absence of regulatory mechanisms that control internalisation of CD4 is unknown, though differences in CD4 levels were previously shown to affect infectivity levels of primary and laboratory-adapted viruses (Kozak et al., 1997). Either extensive endocytosis of receptor-virus complex before initiation of fusion, low expression of CD4 at the cell surface, downstream signalling or unknown interactions could be affected by p56^*lck*^, all of which could influence infection by MCR, and this was therefore investigated further.

HeLa CD4 cells were transduced with a vector expressing p56^*lck*^ and yellow fluorescent protein (YFP) from an internal ribosome entry site (IRES). An immunoblot of p56^*lck*^ and CD4 expression is shown in Supplementary Figure 3, and shows the absence of p56^*lck*^ in untransduced HeLa CD4 and NP2 cells, good expression levels of p56^*lck*^ in transduced cells, and good expression levels of total CD4 across all cells.

The expression of CD4 and CXCR4 at the cell surface in cells with or without added p56^*lck*^ expression was assessed by surface staining through FACS analysis (Supplementary Figure 4 and Supplementary Table 1). The baseline level of anti-CD4 binding to HeLa CD4 cells without the p56^*lck*^ vector (YFP negative) was no different to unstained controls. A small but reproducible increase in levels of anti-CD4 signal was seen for HeLa CD4 cells expressing YFP (and by extension, p56^*lck*^). The significance of this small but persistent increase in median fluorescence intensity level seen on transition from baseline to GFP-negative cells to GFP-positive cells was ascertained by a simple one-tailed rank test. The probability that all experiments give the same ordering of values is p=0.016, meaning that the result is significant at the 95% level. For NP2 cells, p56^*lck*^ /YFP expression made no difference to CD4 cell-surface levels. Additionally, CXCR4 levels for both HeLa CD4 and NP2 cells were not changed by p56^*lck*^ expression, though levels in NP2 cells appeared to be markedly higher.

**Figure 4:**
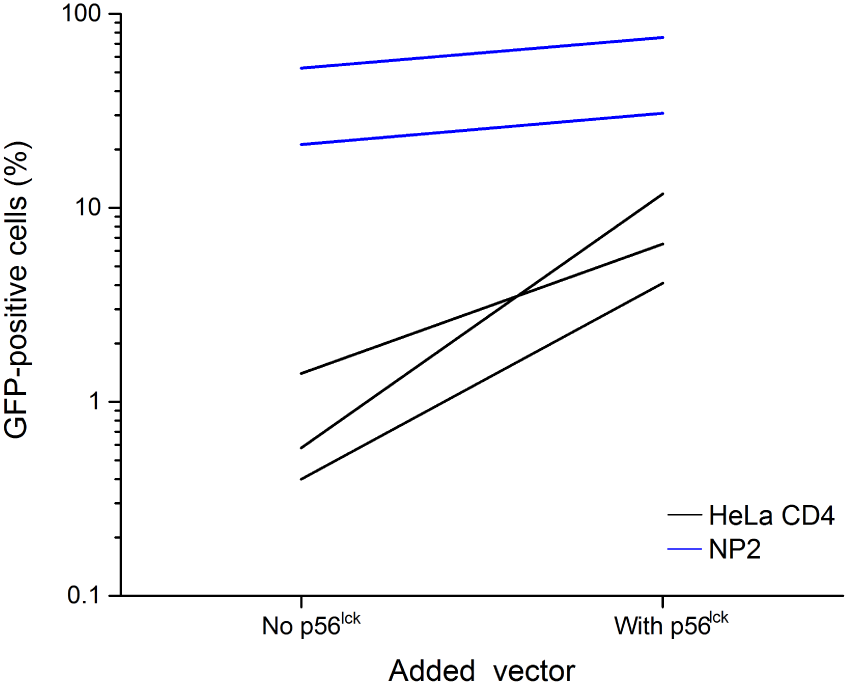
Expression of p56^*lck*^ renders HeLa CD4 cells over ten times more susceptible to infection by MCR Env-pseudotyped virus. Cells were plated and infected with p56^*lck*^ vector at MOI 1-2. After three days, cells were replated and infected by MCR-pseudotyped HIV-1 at MOI 1-2. The extent of infection assayed three days later by FACS and compared to p56^*lck*^-positive and negative cells within the same well, as assessed by YFP fluorescence. Shown are the individual results of three experiments for HeLa CD4 cells, and two for NP2 cells.

A mixed population of HeLa CD4 cells with and without the p56^*lck*^ -YFP vector was then challenged with MCR pseudotyped virus, and the effects of p56^*lck*^ on titres are shown in Figure 4. The two cell lines were plated out together in wells so that a direct comparison could be done under identical conditions, as HeLa CD4 p56^*lck*^ positive cells can be identified by YFP fluorescence. When p56^*lck*^ is expressed in HeLa CD4 cells, titres of MCR ENV-pseudotyped virus increase from undetectable levels (under 1%) to on average around 10-fold higher (Figure 4). Expression of p56^*lck*^ in NP2 cells raised titres by less than two-fold, indicating that the effect of p56^*lck*^ in HeLa CD4 cells is either reduced or absent in NP2 cells (Figure 4).

Taken altogether, these data show that p56^*lck*^ can be overexpressed in HeLa CD4 cells, and that this increases the level of CD4 at the surface of cells. HeLa CD4 cells producing p56^*lck*^ can be infected by MCR Env-pseudotyped virions at levels around 10 times higher than untransduced cells. The change in cell-surface CD4 and increase in titres does not occur in NP2 cells that express p56^*lck*^.

### No enhanced infection level is seen on HeLa CD4 cells expressing p56^*lck*^ compared to HeLa CD4 cells for virions with two non CD4-binding envelope proteins

In order to see if this effect was specific to MCR-mediated viral entry, a comparison of titres of VSV-G, amphotropic envelope and MCR-pseudotyped virions was carried out on HeLa CD4 cells expressing p56^*lck*^, or on the parental p56^*lck*^-null cell line. The expression of p56^*lck*^ in HeLa CD4 cells did not change titres for VSV-G or amphotropic Env-pseudotyped virions (Supplementary Figure 5). For MCR-pseudotyped virions, expression of p56^*lck*^ again increased titres by over 10-fold. While titres for VSV-Gpseudotyped virus are 10-fold higher still, the rescue of MCR-pseudotyped virions was to the same levels as amphotropic Env without p56^*lck*^. This shows that p56^*lck*^ expression can rescue titres of virions pseudotyped with MCR Env, but has no effect on virions pseudotyped with VSV-G or amphotropic Env.

## Discussion

Cells can be artificially engineered to express combinations of retroviral receptors, but expression alone may be insufficient to render these cells permissive to infection by the cognate virus. HeLa CD4 cells are fully permissive to infection by HIV-1 cores pseudotyped with VSV-G, but not to those pseudotyped with an envelope protein derived from a primary isolate, MCR. U87 and NP2 cells, however, are fully permissive to viruses pseudotyped with both envelope proteins. All cell types are clearly capable of supporting retroviral infection. Experiments that compare VSV-G, MCR Env and MCN Env are at best relative, as different virus populations must be used. An attempt was made to recapitulate the blockage using overexpression of Tva800 and Tva950 in HeLa CD4 cells. These are the two forms of receptor for Avian sarcoma leucosis virus which facilitate viral entry through two different routes in the cell, and would permit use of the same virus (Narayan et al., 2003, Gray et al., 2011). However, no difference in viral infection was seen between the two receptors.

As this study shows by qPCR that MCR Env-pseudotyped virions do not enter cells and reverse transcribe at sufficient levels for a productive infection, it was necessary to consider what factor(s) could be responsible for low viral infectivity, including the entry process. Two receptors, CD4 and CXCR4, are necessary for entry via MCR (McKnight et al., 1998). Cloning and sequencing of the CD4 receptor from HeLa CD4 cells showed no non-synonymous mutations compared to the expected coding sequence from *NM*_000616. As overall expression levels of CD4 appeared to be as expected (Supplementary Figure 3 and reference (Oliveira et al., 2010)), we investigated whether a defect in receptor usage could be detected. In order to assess this, a fusion marker, S15-mCherry was used. The data show that MCR Env pseudotyped virions are less likely to fuse with HeLa CD4 cells, and that possibly a reduced absolute number of virions (about 25% and 50% compared to NP2 and U87 cells respectively) bind to HeLa CD4 cells. The qPCR and microscopy data strongly suggest that the block to infection of MCR Env-pseudotyped virions occurs at fusion, and therefore entry.

If successful fusion is not occurring, this could be due to several factors. The virions used to infect HeLa CD4 and NP2 or U87 cells were always the same batch within experiments. There could be a problem with CD4 expression in HeLa CD4 cells, but this is unlikely given it was robustly detected by Western blot of whole cell lysates (Supplementary Figure 3). Interestingly, while these cells are routinely used in CD4-tropic viral titration assays, previous studies showed that for some viruses infectivity on HeLa CD4 cells is correlated with expression level of CD4, including a threshold minimum for NL4-3 Env pseudotyped viruses (Kabat et al., 1994, Kozak et al., 1997, Walter et al., 2005). This study by Kozak et al. reported a similar phenomenon to Lv2 whereby primary patient isolates exhibited different infectivities on U87 and a panel of HeLa CD4 cell clones expressing different levels of CD4 (Kozak et al., 1997). The HeLa CD4 cells used in this and previous studies may well be expressing CD4 at the cell surface that is below this minimum threshold. CD4 is not found at high concentration at the cell surface of non-thymocytes due to rapid endocytosis, and our data with p56^*lck*^ illustrate the importance of cell-surface expression on the observed phenotype. Another possible explanation is that MCR may be binding, but is not able to initiate fusion before the virus-CD4 complex is endocytosed and inactivated by low pH in endosomes. This is not normally a productive entry pathway for HIV (Permanyer et al., 2010).

MCR was derived from a primary isolate, and therefore may be adapted for physiological conditions rather than the tissue culture conditions described in these experiments. It has previously been shown that infectivity of primary isolates is proportional to the level of CD4 expressed on cells, but that this is not a limiting factor for laboratory-adapted strains, and suggested that this is an adaptation to allow these viruses to avoid infections on nonproductive cell types (Kabat et al., 1994, Platt et al., 2000). In vivo, MCR envelope would interact with CD4 on cells of lymphocytic origin, which have regulatory mechanisms to ensure CD4 remains at the cell surface, and not with cells of non-lymphocytic origin. Experiments described here in which titres of MCR pseudotyped virions are increased over 10-fold by expression of p56^*lck*^ support the suggestion that lack of CD4-regulatory mechanisms in HeLa CD4 cells could comprise a component of the block. The addition of p56^*lck*^ restores some part of these mechanisms, and other protein players are highly likely. Neither an increase in the ratio of cell-surface to intracellular CD4, nor a decrease in the rate of endocytosis should affect entry by VSV-G or amphotropic virus (as seen), showing that the effect of p56^*lck*^ is specific to CD4-binding virions.

The block to infection to MCR-pseudotyped viruses in HeLa CD4 cells is only part of a wider phenomenon, and it must be considered why there is no block in NP2 and U87 cells, which, as astrocytoma cells and non-lymphoid in origin, do not normally express endogenous p56^*lck*^. However, it has been suggested that p56^*lck*^ is expressed in rat and mouse brains (Omri et al., 1996, Van Tan et al., 1996). This serves to highlight the difference between cells of different lineages and origins. If the difference between NP2, U87 and HeLa CD4 were to be probed further, it would be interesting to widen the screen well beyond p56^*lck*^, and to look at expression profiles across HeLa, NP2, U87 and lymphocytes of other proteins known to interact with CD4 as other factors may be responsible for the regulation of CD4 (or lack thereof) in other cell types. The titration data for MCN Env and NL4-3 Env pseudotyped virions show that the block to infection is not unique to MCR Env, and further CD4/CXCR4-tropic envelopes could also be investigated.

It has been suggested that the difference between HeLa CD4 and NP2 cells is due to an intracellular restriction factor present in a specific compartment of HeLa CD4 cells that inhibits viral cores similarly to Trim5*α* or Fv1 (Stremlau et al., 2004, Best et al., 1996, Schmitz et al., 2004), and that rerouting virus around this compartment (e.g. by VSV-G) enables the virus to avoid this block. For a late entry block (e.g. Fv1n) it can be ascertained that fusion and entry are occurring by comparing levels of both early and late reverse transcription products (i.e. for N-(unrestricted) and B-(restricted) tropic MLV (Jolicoeur and Rassart, 1980)). For an early block, (e.g. Trim5*α*) levels of early products of restricted and unrestricted virus are not comparable as the block is pre-reverse transcription (Stremlau et al., 2004). However, the saturable nature of Trim5*α* (and Fv1) overwhelmingly indicates that a post-entry, cytosolic restriction factor is involved, rather than a block at the level of fusion and entry, whereas no abrogation was achieved for MCR-pseudotyped virus on HeLa CD4 cells (Supplementary Figure 2, see (Dodding et al., 2005, Hatziioannou et al., 2003) for comparison). The block could be either nonsaturable in nature, or expressed at too high a concentration to saturate. In HeLa CD4 cells infected with MCR-pseudotyped virus, levels of reverse transcription products do not accrue to those comparable to a successful infection and it is less likely that the block is definitively caused by a specific restriction factor, or through passage via a particular cell compartment, as there is no indication that the infection would otherwise be successful.

These experiments were carried out in order to investigate an aspect of Lv2 as proposed by previous studies. Instead of independently creating an exact replica of previous studies, we sought instead to use different methods to shed light from an alternative angle. Where experiments from prior studies were repeated, the data shown here were similar to those previously. To take Figure 2 as an example, Schmitz et al. showed ratios of strong-stop products produced from HeLa CD4 cells and compared them to the levels from U87 cells for both MCR- and MCN-pseudotyped viruses, showing the data as a fold-restriction ratio (Schmitz et al., 2004). The fold-difference (i.e. the ratio of transcripts) in U87 cells at all timepoints for MCR Env-pseudotyped virions is between 7 to 25fold higher than in HeLa CD4 cells. Those data are in agreement with these presented here, as Figure 2 shows a strong-stop products ratio of approximately 5-fold of unrestricted cells compared to HeLa CD4. Again, data from a similar (though non-quantitative) experiment are shown in McKnight et al. (McKnight et al., 2001), where the two initial primary isolates, prCBL23 and CBL23, are compared. Although the levels of reverse transcripts for the two isolates are the same when considered on each cell type individually, between cell types the levels are very different (with much higher levels produced from U87 cells than the restrictive HeLa CD4 or HOS cells). Those data are again commensurate with Figure 2 of this study.

Data from this study can also be compared to previous studies with different methodologies. In Reuter et al. an experimental technique that discriminates between fusion and endocytosis was used to investigate the block to MCR pseudotyped virions, known as the BlaM-vpr assay, from which the results can be compared to those in Figure 3 (Cavrois et al., 2002, Reuter et al., 2005). Briefly, Reuter et al. reported that entry via restricted Env proteins, which included Env35 (or MCR), was between 2.5 to 3-fold less efficient compared to entry via unrestricted Env proteins. This is in concordance with the results in Figure 3, where a level of 4 to 5-fold less fusion was observed in restricted HeLa CD4 cells compared to unrestricted NP2 or U87 cells. Again, the data are in agreement, though the subsequent conclusions drawn may differ.

In experiments detailed in this study we have studied a phenomenon carefully by standard and novel techniques, and report that while there is a block to infection of MCR-pseudotyped virions in HeLa CD4 cells, this is at entry and could be due to phenomena other than a restriction factor (Goff, 2004, Sorin and Kalpana, 2006). Experiments described here confirm the original phenotype of MCR Env and the results are concordant with those from previous papers. Some cells can be infected by MCR Env pseudotyped virus, but other cells expressing the same combination of receptor and co-receptor are not, but this artificial combination of cells and receptors could highlight an artefact of cell biology, and CD4 interaction with the Env of primary isolates (Kabat et al., 1994). Virion entry levels are not commensurate with a successful infection, and fusion of MCR with HeLa CD4 cells is reduced compared to NP2 or U87 cells. The block to infection, therefore, occurs at viral entry, and can be modulated by cellular factors that interact with CD4. A plausible explanation of the primary cause of this phenomenon is differences in cell biology between NP2, U87 and HeLa CD4 cells with inhibition of the viral lifecycle at the stages of fusion and entry.

## Additional Information

### Acknowledgements

ERG thanks Melvyn Yap for technical assistance and discussions, Laura Hilditch for randomising the microscopy images and for comments on the manuscript; Daniel Ashley Richards, Petra Mlcochova and Doug King for access to facilities. This work was supported by a Medical Research Council studentship to ERG and conducted in the laboratory of Dr Jonathan Stoye, with additional support from EPSRC EP/K031953/1 i-sense: Early Warning Sensing Systems for Infectious Disease.

